# Common gene expression signatures in Parkinson’s disease are driven by changes in cell composition

**DOI:** 10.1101/778910

**Authors:** Gonzalo S. Nido, Fiona Dick, Lilah Toker, Kjell Petersen, Guido Alves, Ole-Bjørn Tysnes, Inge Jonassen, Kristoffer Haugarvoll, Charalampos Tzoulis

## Abstract

**Background:** The etiology of Parkinson’s disease (PD) is largely unknown. Genome-wide transcriptomic studies in bulk brain tissue have identified several molecular signatures associated with the disease. While these studies have the potential to shed light into the pathogenesis of PD, they are also limited by two major confounders: RNA post mortem degradation and heterogeneous cell type composition of bulk tissue samples. We performed RNA sequencing following ribosomal RNA depletion in the prefrontal cortex of 49 individuals from two independent case-control cohorts. Using cell-type specific markers, we estimated the cell-type composition for each sample and included this in our analysis models to compensate for the variation in cell-type proportions.

**Results:** Ribosomal RNA depletion results in substantially more even transcript coverage, compared to poly(A) capture, in post mortem tissue. Moreover, we show that cell-type composition is a major confounder of differential gene expression analysis in the PD brain. Correcting for cell-type proportions attenuates numerous transcriptomic signatures that have been previously associated with PD, including vesicle trafficking, synaptic transmission, immune and mitochondrial function. Conversely, pathways related to endoplasmic reticulum, lipid oxidation and unfolded protein response are strengthened and surface as the top differential gene expression signatures in the PD prefrontal cortex.

**Conclusions:** Differential gene expression signatures in PD bulk brain tissue are significantly confounded by underlying differences in cell-type composition. Modeling cell-type heterogeneity is crucial in order to unveil transcriptomic signatures that represent regulatory changes in the PD brain and are, therefore, more likely to be associated with underlying disease mechanisms.

## Introduction

Parkinson’s disease (PD) is the second most prevalent neurodegenerative disorder, affecting ∼1.8 % of the population above 65 years [1]. PD is a complex disorder, caused by a combination of genetic and environmental factors, but the molecular mechanisms underlying its etiology remain largely unaccounted for. Genome-wide transcriptomic studies can identify expression signatures associated with PD. While not able to establish causality, these studies hold the potential to highlight important biological mechanisms, some of which may be exploited as targets for therapeutic modulation.

A recent systematic review identified 33 original genome-wide transcriptomic studies in the PD brain, of which 6 were performed on laser microdissected neurons from the *substantia nigra pars compacta* (SNc) and the remaining in bulk tissue from various brain regions [2]. These studies show surprisingly low replicability at the level of individual genes, however, and only partial concordance for pathways. The most consistent alterations have been found in pathways related to energy metabolism/mitochondrial function and protein degradation, followed by synaptic transmission, vesicle trafficking, lysosome/autophagy and neuroinflammation [2]. While these processes commonly show differential expression signatures in PD, it remains unknown whether this is because they truly reflect the biology of PD or due to systematic bias and confounding factors. Two major sources of bias for transcriptomic studies in the human brain are the *post mortem* degradation of RNA and the highly heterogeneous cell type composition of bulk tissue samples.

RNA degradation of variable extent occurs in *post mortem* tissue. To further complicate the picture, it has been shown that different cell types exhibit different degrees of susceptibility to RNA degradation [3], potentially confounding differences in cellular composition with differences in RNA quality. Access to high-quality brain tissue is generally limited, and thus an optimal choice of experimental platforms becomes paramount to maximize sensitivity. While RNA microarrays are being gradually superseded by RNA-seq technology, only 3 out of the 33 studies identified by an up-to-date review [2] used RNA-seq, and all of them employed poly(A) capture, a widely used protocol to restrict the analysis to mature mRNA [4–6]. However, this library preparation method only picks up RNA fragments with a poly-A tail, introducing substantial bias in low quality RNA samples [7–10]. A well-established approach to mitigate this limitation, is whole RNA-seq following ribosomal RNA (rRNA) depletion [11]. To our knowledge there are no genome-wide transcriptomic studies on PD brain employing rRNA depletion methods to date.

Systematic differences in sample cell composition represent another important confounding factor. These typically originate from two sources: biological differences (e.g. secondary to neurodegeneration) and technical variation in sample dissection and preparation. Brain areas affected by neurodegeneration are characterized by neuronal loss and gliosis, resulting in a systematically increased glia-to-neurons ratio in patients. This confounder is strongest in areas with severe changes, such as the SNc, but is also present to a variable degree in less affected areas, such as the neocortex. In addition, technical sources of variation due to sampling may affect any brain region and cause an uneven distribution of gray and white matter, resulting in a variable fraction of oligodendrocytes. Thus, transcriptional signatures associated with PD in bulk brain tissue may reflect changes in cellular composition rather than disease-specific transcriptional modulation. This observation has already been put forward using neurodegenerative mouse models and re-analysis of human brain transcriptomic data [12]. Heterogeneous cell composition is, hence, a major confounder that needs to be considered and appropriately addressed in transcriptomic studies in bulk brain samples.

We report the first genome-wide transcriptomic study in the PD brain employing RNA-seq following rRNA depletion. We show that this approach is able to substantially mitigate the bias of *post mortem* degradation, resulting in substantially better transcript coverage compared to poly(A) capture. Moreover, by estimating the relative cell type proportion in our samples, we confirm that cellular composition is a major source of variation in bulk tissue data, confounding the differential gene expression profile even in the less affected prefrontal cortex. By incorporating the estimated cell type proportions into our analysis models, we were able to unveil transcriptomic signatures which are more likely to be associated with the underlying disease mechanisms.

## Results

### Ribosomal RNA depletion is superior to poly(A) selection in *post mortem* brain

We carried out RNA-seq following rRNA depletion in fresh-frozen prefrontal cortex (Brodmann area 9) from a total of 49 individuals from two independent cohorts: the Norwegian ParkWest study (PW, n = 29) [13] and the Netherlands Brain Bank (NBB, n = 20). A detailed description of our cohorts is provided in the methods. Comparison of our data to a published poly(A) capture dataset of similar characteristics [4] revealed important differences of mapping coverage. Mapping efficiency was slightly higher in the poly(A) dataset (median = 0.976, range = 0.971-0.980) compared to the rRNA-depleted datasets (PW: median = 0.952, range = 0.940-0.962; NBB: median = 0.959, range = 0.947-0.965). The counts per million (CPMs) of rRNA regions, as defined by Ensembl release 75, was very low in all samples (PW: median = 3,099, range = 1,047-7,071; NBB: median = 1,583, range = 1,129-5,024) and, as expected, significantly lower in the rRNA-depleted cohorts than in the poly(A) capture (poly(A) cohort: median = 40,058, range = 10,701-95,183) (S1 Fig).

In both datasets, the RNA integrity number (RIN) was positively correlated with mapping efficiency to mRNA regions, but not to intergenic and/or intronic regions (S2 Fig). Despite having higher mean RIN values, the poly(A) capture cohort showed a marked unevenness of transcript body coverage compared to the rRNA depletion samples (Fig 1A). The median coefficient of variation in coverage was significantly lower in the rRNA-depleted samples and the 5’- and 3’-ends of the transcripts showed substantially better coverage compared to the poly(A) sample (Fig 1B). Moreover, in the rRNA-depleted samples both the 3’- and 5’-end coverage loss showed a significant inverse correlation with the RIN values. In contrast, RIN showed no correlation with the 5’-bias and a positive correlation with the 3’-bias in the poly(A) capture samples (Fig 1C). Thus, rRNA depletion results in substantially better and more even coverage of the transcriptome in *post mortem* brain tissue, providing a better alternative to poly(A) capture and minimizing the prospect of transcript quantification biases downstream.

**Fig 1.**
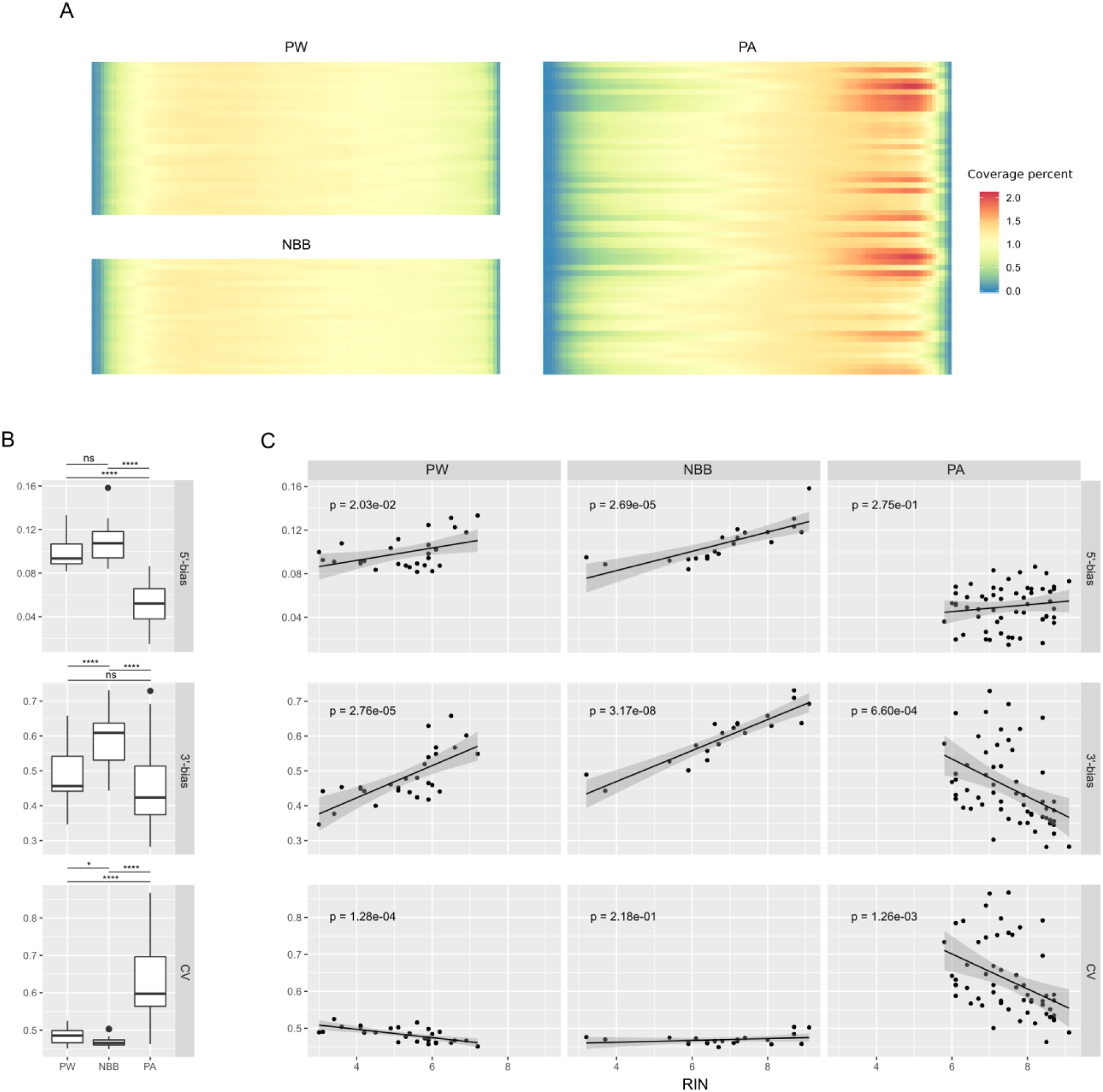
Transcript coverage profiles with rRNA depletion compared to poly(A) capture. (A) Heatmaps of transcript coverage in our two cohorts (PW, NBB) and a poly(A) capture sample (PA). The y-axis shows samples sorted by RIN (top: lowest RIN; bottom highest RIN). The x-axis represents the transcript body percentiles (5’ to 3’). The shading for a given row represents the sample-normalized coverage averaged across all transcripts. (B) Boxplots for different coverage quality metrics: median 5’-bias, median 3’-bias and median coefficient of variation (CV) for each cohort. The bias metric is calculated by Picard tools on the 1000 most highly expressed transcripts and corresponds to the mean coverage of the 3’(or 5’)-most 100 bases divided by the mean coverage of the whole transcript (asterisks indicate significance at (*) p > 0.05, (**) p ≤ 0.01, (***) p ≤ 0.001, (****) p ≤ 0.0001, Wilcoxon test). The same metrics are expanded in (C), with sample scatterplots showing RIN values against the coverage quality metrics. Linear regression trends are indicated with black lines. P-values for the F-statistic of the linear model are also shown in the panels. Panels are organized in columns (cohorts) and quality metrics (rows). CV = coefficient of variation; PW = ParkWest cohort; NBB = Netherlands Brain Bank cohort; PA = poly(A) cohort

### Cell composition is a major confounder of gene expression in bulk brain samples

The observed gene expression profiles in bulk brain tissue can be dramatically influenced by differences in cellular composition. Such differences can be a result of variation in gray/white matter ratios introduced during tissue extraction, inter-subject variability or represent disease related alterations [14–16]. To study the contribution of various technical and biological sources of variation in our dataset we first estimated marker gene profiles (MGPs) for the major classes of cortical cell types (astrocytes, microglia, oligodendrocytes, and neurons) in our samples by summarizing the expression of the cell type-specific marker genes as previously described [15]. Next, we examined the Pearson’s correlation between potential sources of biological variation in our data, including technical and demographic factors (RIN, *post-mortem* interval (PMI), sex, age, and disease status) and MGPs. As expected, MGPs for neuronal cell types were significantly anticorrelated with the other main cortical cell types in both cohorts (p < 0.05, Fig 2A). In agreement with previous studies [3, 17], cellular estimates were also confounded with RNA quality. In both cohorts RIN was significantly correlated with neuronal (positive correlation) and astrocyte (negative correlation) MGPs. Significant negative correlation of RIN with microglia MGPs was observed in the NBB cohort (Fig 2A). Most concerning was the detection of a significant association between the oligodendrocyte MGP and the disease status in the PW cohort (Fig 2A). The main axis of variation in gene expression (which explained 44% and 45% of the total variance in PW and NBB, respectively) was significantly correlated with RNA quality and cellular composition in both cohorts (Fig 2B), singling out RNA quality and cellular composition as the main drivers of transcriptional change in bulk brain tissue.

**Fig 2.**
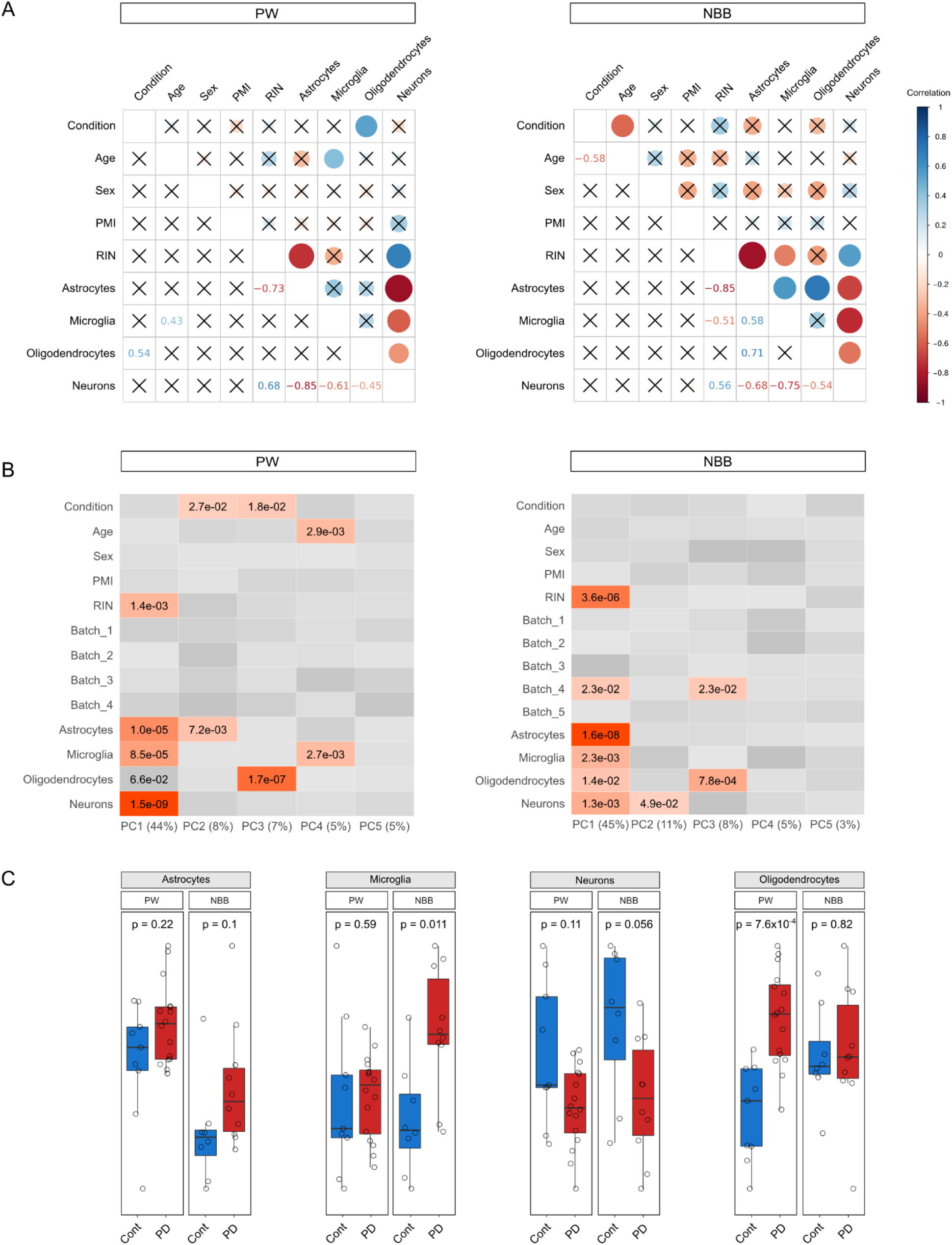
Analysis of sample covariates. (A) Pearson correlation coefficients for each pair of variables are shown in correlograms. Sizes of the circles in the upper triangular of the correlograms are proportional to the Pearson correlation coefficient, with color indicating positive (blue) or negative (red) coefficients. The precise values for the Pearson coefficients are indicated in the lower triangular. Non-significant pairwise correlations (p ≥ 0.05) are represented with a cross. (B) Heatmaps showing the association between the sample variables with the first 5 principal components of the gene expression. Only significant p-values (p < 0.05) are represented (linear regression F-test). (C) Cell type estimates based on MGPs for the main cortical cell types controlling for all the experimental variables except disease status (i.e. sex, age, PMI, RIN, and sequencing batch). P-values calculated with Wilcoxon tests. PW = ParkWest cohort; NBB = Netherlands Brain Bank cohort

We next looked for differences in cellular proportions between PD and controls adjusting for the known experimental covariates (see methods). In the NBB cohort, PD subjects exhibited a significant increase in the microglia MGP (p = 0.015, Wilcoxon test), while a significant increase in the oligodendrocyte MGP (p = 5.5×10^−3^, Wilcoxon test) was observed in PD subjects from the PW cohort. In both cohorts, these changes were accompanied by a non-significant decrease in neuronal MGPs (Fig 2C).

Since the oligodendrocyte and microglia MGPs were significantly associated with the disease status in at least one of the cohorts, we re-estimated group differences in astrocyte and neuronal MGPs, adjusting for the significant MGPs, in addition to the original covariates. This analysis showed no significant differences between the groups (S3 Fig). Therefore, only MGPs of oligodendrocytes and microglia were included in the statistical model of differential expression analysis.

### Differential gene expression

Differential gene expression analysis of a total of ∼31,000 pre-filtered genes was carried out using experimental covariates (sex, age, PMI, RIN, and sequencing batch) with or without oligodendrocyte and microglia MGPs. In the PW cohort, 595 genes were defined as differentially expressed (Benjamini-Hochberg FDR < 0.05) without adjusting for cell type composition. Inclusion of oligodendrocyte and microglia MGPs in the model decreased the number of differentially expressed genes to a total of 220. In total, 74 genes remained significant both with and without adjustment for cell type composition. No transcripts with FDR < 0.05 were identified in the NBB cohort, irrespective of adjustment for cell type composition. A list with the nominally significant genes overlapping between the two cohorts is provided in S1 File. Comprehensive results of differential expression analysis are available in S2 File.

### Functional enrichment

Functional enrichment analysis of the differential gene expression results without MGP adjustment indicated 476 significantly enriched (Benjamini-Hochberg FDR < 0.05) pathways in PW (107 up-regulated and 369 down-regulated) and 992 in NBB (421 up-regulated and 571 down-regulated). MGP adjustment reduced the number of significant pathways to 89 in PW (35 up-regulated and 54 down-regulated) and 248 in NBB (115 up-regulated and 133 down-regulated). Of these, 34 pathways replicated across the two cohorts. Concordant pathways comprised protein folding, ER-related processes and lipid oxidation (Fig 3). The complete results are provided in S3 File.

**Fig 3.**
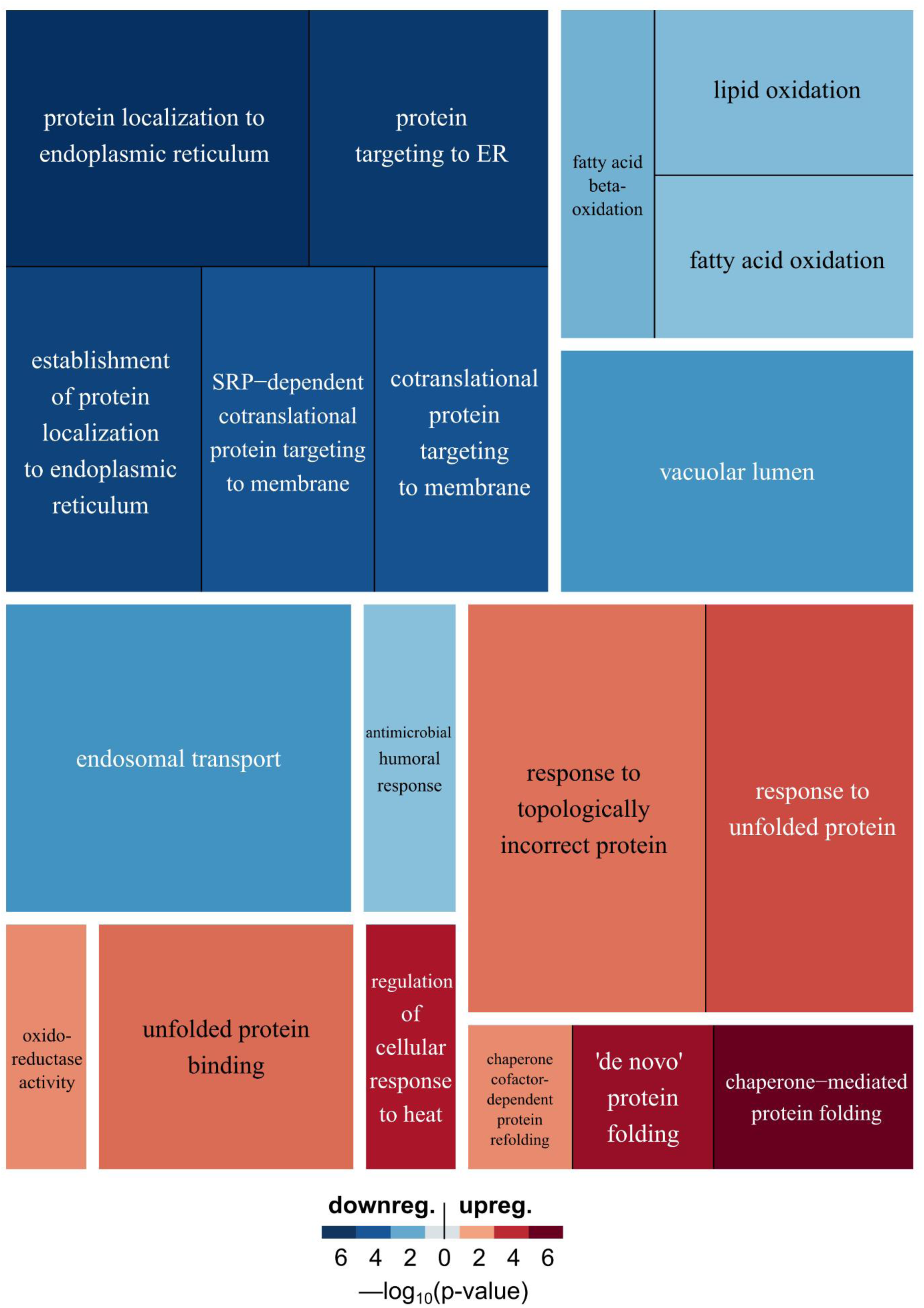
Functional enrichment. The treemap shows the concordant enriched pathways between PW and NBB accounting for experimental covariates and MGPs (same direction of gene expression change and Benjamini-Hochberg FDR < 0.05). Pathways are grouped with a white border if their gene overlap is above 0.5 (Szymkiewicz–Simpson coefficient). Darker shades of red/blue represent lower enrichment p-values for up-/down-regulated pathways. Sizes of the rectangles are proportional to pathway sizes

As expected, scoring each pathway according to the change in p-value when accounting for cellularity, revealed a marked downplay of the relevant cell-type-specific functions (Table 1). In the PW cohort, which was characterized by a skewed oligodendrocytes/neurons proportion, the function with the largest attenuation (i.e. increase in p-value) was seen for up-regulation of myelination and other oligodendrocyte related functions and for down-regulation of neuronal pathways. For NBB, correcting for cell-composition resulted in attenuation of immunity and neuronal pathways, consistent with the unbalanced microglial/neuronal proportions seen in that cohort (Table 1). Strikingly, pathways linked to mitochondrial respiration, including respiratory complex I, were among the down-regulated processes that lost statistical significance when controlling for cellularity. The attenuation of the mitochondrial signal was observed in both cohorts. Conversely, up-regulation of protein folding-related pathways gained significance in both cohorts (Table 2 and Fig 3). Complete results are provided in S4 File.

**Table 1.** Loss of significance in enriched pathways.

| PW |  |  |  |
| --- | --- | --- | --- |
| Up-regulated |  | Down-regulated |  |
| *Pathway* | *Delta* | *Pathway* | *Delta* |
| myelination | -8.85 | regulation of synaptic vesicle exocytosis | -9.63 |
| ensheathment of neurons | -8.67 | intrinsic component of synaptic membrane | -9.60 |
| axon ensheathment | -8.67 | regulation of synaptic vesicle cycle | -9.58 |
| detection of chemical stimulus involved in sens... | -7.95 | positive regulation of synaptic transmission | -9.48 |
| oligodendrocyte differentiation | -6.94 | Schaffer collateral - CA1 synapse | -9.46 |
| oligodendrocyte development | -6.58 | regulation of synaptic plasticity | -9.44 |
| apical junction complex | -5.57 | regulation of neurotransmitter secretion | -9.41 |
| glial cell development | -5.54 | presynaptic membrane | -9.27 |
| glial cell differentiation | -5.52 | regulation of synaptic vesicle transport | -9.19 |
| tight junction | -4.16 | protein transport within lipid bilayer | -9.11 |

| NBB |  |  |  |
| --- | --- | --- | --- |
| Up-regulated |  | Down-regulated |  |
| *Pathway* | *Delta* | *Pathway* | *Delta* |
| activation of innate immune response | -8.54 | ribonucleoside monophosphate metabolic process | -9.09 |
| regulation of leukocyte proliferation | -8.11 | purine nucleoside triphosphate metabolic process | -9.03 |
| regulation of lymphocyte proliferation | -8.01 | mitochondrial membrane part | -8.99 |
| regulation of mononuclear cell proliferation | -7.79 | ATP metabolic process | -8.92 |
| innate immune response-activating signal transd... | -7.43 | regulation of synaptic vesicle exocytosis | -8.87 |
| regulation of adaptive immune response | -7.16 | inner mitochondrial membrane protein complex | -8.82 |
| response to interferon-gamma | -7.15 | purine ribonucleoside triphosphate metabolic proc... | -8.77 |
| adaptive immune response based on somatic rec... | -7.07 | respiratory chain | -8.70 |
| blood microparticle | -6.75 | regulation of synaptic vesicle transport | -8.68 |
| regulation of T cell proliferation | -6.73 | cellular respiration | -8.43 |

Tables representing the top 10 pathways with the lowest delta for up- and down-regulated pathways for both cohorts. The delta value represents the change in the enrichment $-\log_{10}(\text{p-value})$ between the results with and without MGP adjustment (negative values of delta imply a loss of significance when accounting for cellularity). Complete results are provided in S4 File.

**Table 2.** Gain of significance in enriched pathways.

| PW |  |  |  |
| --- | --- | --- | --- |
| Up-regulated |  | Down-regulated |  |
| *Pathway* | *Delta* | *Pathway* | *Delta* |
| protein folding | 5.77 | DNA packaging complex | 3.76 |
| 'de novo' protein folding | 5.73 | basement membrane | 3.47 |
| unfolded protein binding | 5.54 | positive regulation of epithelial cell proliferation | 2.52 |
| chaperone-mediated protein folding | 5.32 | negative regulation of gliogenesis | 2.49 |
| 'de novo' posttranslational protein folding | 4.68 | fatty acid beta-oxidation | 2.30 |
| heat shock protein binding | 4.34 | nucleosome | 2.18 |
| response to unfolded protein | 4.10 | glomerulus development | 2.16 |
| response to topologically incorrect protein | 3.53 | aorta development | 2.06 |
| oxidoreductase activity, acting on paired... | 2.74 | endothelium development | 1.95 |

| NBB |  |  |  |
| --- | --- | --- | --- |
| Up-regulated |  | Down-regulated |  |
| *Pathway* | *Delta* | *Pathway* | *Delta* |
| positive regulation of cardiac muscle tissue dev... | 1.99 | tertiary granule | 5.22 |
| regulation of smooth muscle cell differentiation | 1.98 | ficolin-1-rich granule membrane | 5.00 |
| negative regulation of protein serine/threonine kin... | 1.98 | regulation of myeloid leukocyte mediated immunity | 4.55 |
| hormone-mediated signaling pathway | 1.95 | regulation of leukocyte degranulation | 4.34 |
| lung alveolus development | 1.71 | specific granule | 4.22 |
| positive regulation of striated muscle tissue dev... | 1.69 | ficolin-1-rich granule | 4.15 |
| positive regulation of muscle organ development | 1.69 | tertiary granule membrane | 3.57 |
| positive regulation of muscle tissue development | 1.62 | regulation of mast cell activation | 3.52 |
| negative regulation of MAP kinase activity | 1.48 | vacuolar lumen | 3.05 |
| regulation of cardiac muscle cell differentiation | 1.48 | regulation of mast cell degranulation | 3.05 |

Tables representing the top 10 pathways with the highest delta for up- and down-regulated pathways for both cohorts. The delta value represents the change in the enrichment $-\log_{10}(p\text{-value})$ between the results with and without MGP adjustment (positive values imply an increase in p-value when accounting for cellularity). Complete results are provided in S4 File

## Discussion

We present the first genome-wide transcriptomic study in the PD brain employing whole RNA-seq after rRNA depletion. Our findings show that PD-associated differential gene expression signatures in bulk brain tissue are influenced to a great extent by the underlying differences in cell-type composition of the samples. Modeling cell-type heterogeneity allowed us to highlight transcriptional signatures that are likely to represent aberrant gene expression within the cells of the PD brain, rather than changes in cell composition.

Our results suggest that rRNA depletion is superior to the more commonly used poly(A) capture approach and allows for a more accurate mapping and quantification of the transcriptome in *post mortem* brain tissue. The rRNA depletion method provides substantially higher evenness of coverage and effectively mitigates the 3’- and 5’-end coverage bias associated with poly(A) capture [7–10]. Ultimately, the unevenness of coverage will influence transcript quantification, affecting the sensitivity of the differential expression estimates.

Our study supports the notion that cell composition can be a major confounder in bulk brain tissue transcriptomics. The observed expression profiles in both cohorts were driven predominantly by a combination of technical factors associated with RNA quality, and differences in cellular composition between PD and controls. This difference was primarily due to oligodendrocytes in PW and microglia in NBB. Since oligodendrocyte proliferation is not a pathological feature of PD, it is plausible that the difference in oligodendrocyte MGPs in PW was due to technical variation in gray/white matter content introduced during tissue sampling. Microglial infiltration does occur in affected areas of the PD brain [18]. It is noteworthy, however, that increased microglial MGP was only observed in one of the cohorts (NBB), highlighting the biological heterogeneity of PD. Accounting for relative cell proportions reduced the number of differentially expressed genes and attenuated the calculated enrichment of cell-type-specific pathways between PD and controls. In the PW cohort, this alleviated a substantial false positive signature of oligodendrocyte genes presumably caused by skewed grey/white matter sampling bias. Similar sampling bias could be responsible for oligodendroglia-specific functions appointed to PD brain in previous transcriptomic studies [6].

Intriguingly, accounting for cellular proportions downplayed several of the transcriptomic signatures that have been previously associated with PD. For instance, the signal from vesicle trafficking- and synaptic transmission-related processes [19–24] was significantly attenuated in both cohorts, suggesting that the signal was primarily driven by changes in neuronal proportions between PD and controls, rather than modulation of these pathways within neurons. Moreover, we observed an attenuation in the down-regulation of mitochondrial pathways, including the respiratory chain and oxidative phosphorylation, which are among the most consistent transcriptomic signatures in PD [2,4,19,21,25–28]. The loss of transcriptional signal in these pathways is intriguing, because there is compelling evidence that decreased complex I protein levels occur in PD neurons [29]. Our results suggest that the previously reported transcriptional down-regulation of the respiratory chain is at least partly driven by altered cellular composition (due to decreased number of neurons which highly express these genes) and may therefore not be the sole mechanism by which neuronal complex I deficiency occurs in PD. Indeed, it has been suggested that complex I deficiency in PD may be mediated by proteolytic degradation by the LON-ClpP protease system, rather than transcriptional regulation [30].

Similar changes in the cell-composition of the brain are seen in all neurodegenerative diseases, including PD, Alzheimer disease, ALS and Huntington disease. Interestingly, common and overlapping transcriptional signatures have been reported across these neurodegenerative diseases, including mitochondrial, neuronal-specific, and immunity-related pathways [31, 32]. Our findings suggest that these common transcriptional signatures of neurodegeneration may be largely confounded by a common pattern of altered cellularity, involving neuronal loss and glial proliferation, rather reflect biological processes of causal nature.

Correcting for cell-type composition in our samples highlighted processes related to the endoplasmic reticulum, unfolded protein response and lipid/fatty acid oxidation as the top differential gene expression signatures in the PD prefrontal cortex. Unfolded protein response is indeed one of the most consistently reported transcriptomic signatures in PD [2,4,21,25,33,34]. Moreover, endoplasmic reticulum stress and aberrant proteostasis have been associated with the accumulation of misfolded proteins, including α-synuclein, in both *in vitro* studies and animal models of PD [35]. While less is known regarding the role of lipid metabolism in PD, evidence of aberrant fatty acid oxidation has been found by metabolomic studies in serum [36] and urine [37] of patients. Our results corroborate these findings and indicate that aberrant fatty acid metabolism occurs in the PD prefrontal cortex.

Based on our findings, we advocate that modeling cell-type heterogeneity is crucial in order to unveil transcriptomic signatures reflecting regulatory changes in the PD brain. It is, however, important noting that modeling of cellular estimates cannot completely mitigate the cell-composition bias in bulk tissue. Moreover, cell-type correction complicates identification of transcriptional events co-occurring with changes in cellular composition. Single-cell or cell-sorting based methods will be key to overcoming this limitation and deciphering transcriptomic signatures directly associated with underlying disease mechanisms in PD.

## Materials and methods

### Subject cohorts

All experiments were conducted in fresh-frozen prefrontal cortex (Brodmann area 9) from a total of 49 individuals from two independent cohorts. The first cohort (n = 29) comprised individuals with idiopathic PD (n = 18) from the Park-West study (PW), a prospective population-based cohort which has been described in detail [38] and neurologically healthy controls (Ctrl, n = 11) from our brain bank for aging and neurodegeneration. Whole-exome sequencing had been performed on all patients [39] and known/predicted pathogenic mutations in genes implicated in Mendelian PD and other monogenic neurological disorders had been excluded. None of the study participants had clinical signs of mitochondrial disease or use of medication known to influence mitochondrial function (S5 File). Controls had no known neurological disease and were matched for age and gender. The second cohort comprised samples from 21 individuals from the Netherlands Brain Bank (NBB) including idiopathic PD (n = 10) and demographically matched neurologically healthy controls (n = 11). Individuals with PD fulfilled the National Institute of Neurological Disorders and Stroke [40] and the UK Parkinson’s disease Society Brain Bank [41] diagnostic criteria for the disease at their final visit.

To investigate the effect of rRNA depletion compared to the prevailing poly(A) capture method, we re-analyzed an RNA-seq dataset from a previous publication which employed a poly(A) tail selection kit on post-mortem tissue of the same brain area and same disease (poly(A) cohort, n = 29 PD samples, n = 44 neurologically healthy controls, all males; GEO: GSE68719) [4].

Informed consent was available from all individuals. Ethical permission for these studies was obtained from our regional ethics committee (REK 2017/2082, 2010/1700, 131.04).

### Tissue collection and neuropathology

Brains were collected at autopsy and split sagittaly along the *corpus callosum*. One hemisphere was fixed whole in formaldehyde and the other coronally sectioned and snap-frozen in liquid nitrogen. All samples were collected using a standard technique and fixation time of ∼ 2 weeks. There was no significant difference in PMI (independent t-test, PW cohort p = 0.16; NBB cohort p = 0.92), age (independent t-test, PW cohort p = 0.18; NBB cohort p = 0.074) or gender (independent t-test, PW cohort p = 0.94; NBB cohort p = 0.53) between PD subjects and controls. Subject demographics and tissue availability are provided in S5 File. Routine neuropathological examination including immunohistochemistry for α-synuclein, tau and beta amyloid was performed on all brains. All cases showed neuropathological changes consistent with PD including degeneration of the dopaminergic neurons of the SNc in the presence of Lewy pathology. Controls had no pathological evidence of neurodegeneration.

### RNA sequencing

Total RNA was extracted from prefrontal cortex tissue homogenate for all samples using RNeasy plus mini kit (Qiagen) with on-column DNase treatment according to manufacturer’s protocol. Final elution was made in 65 µl of dH2O. The concentration and integrity of the total RNA was estimated by Ribogreen assay (Thermo Fisher Scientific), and Fragment Analyzer (Advanced Analytical), respectively and 500 ng of total RNA was used for downstream RNA-seq applications. First, rRNA was removed using Ribo-Zero™ Gold (Epidemiology) kit (Illumina, San Diego, CA) using manufacturer’s recommended protocol. Immediately after the rRNA removal the RNA was fragmented and primed for the first strand synthesis using the NEBNext First Strand synthesis module (New England BioLabs Inc., Ipswich, MA). Directional second strand synthesis was performed using NEBNExt Ultra Directional second strand synthesis kit. Following this the samples were taken into standard library preparation protocol using NEBNext® DNA Library Prep Master Mix Set for Illumina® with slight modifications. Briefly, end-repair was done followed by poly(A) addition and custom adapter ligation. Post-ligated materials were individually barcoded with unique in-house Genomic Services Lab (GSL) primers and amplified through 12 cycles of PCR. Library quantity was assessed by Picogreen Assay (Thermo Fisher Scientific), and the library quality was estimated by utilizing a DNA High Sense chip on a Caliper Gx (Perkin Elmer). Accurate quantification of the final libraries for sequencing applications was determined using the qPCR-based KAPA Biosystems Library Quantification kit (Kapa Biosystems, Inc.). Each library was diluted to a final concentration of 12.5 nM and pooled equimolar prior to clustering. 125 bp Paired-End (PE) sequencing was performed on an Illumina HiSeq2500 sequencer (Illumina, Inc.). RNA quality, as measured by the RNA integrity number (RIN), varied across samples (mean = 5.3, range = 3.0-7.2 for PW; mean = 6.8, range = 3.2-9.1 for NBB, S5 File), although the difference between conditions did not reach statistical significance in any of the cohorts (t-test P = 0.72 and 0.90 for PW and NBB cohorts, respectively).

### Data quality control

FASTQ files were trimmed using Trimmomatic version 0.36 [42] to remove potential Illumina adapters and low quality bases with the following parameters: ILLUMINACLIP:truseq.fa:2:30:10 LEADING:3 TRAILING:3 SLIDINGWINDOW:4:15. FASTQ files were assessed using fastQC version 0.11.5 [43] prior and following trimming. For an in-depth quality assessment, we mapped the trimmed reads using HISAT2 version 2.1.0 [44] against the hg19 human reference genome (using --rna-strandness RF option) preserving lane-specific information. To discard potential lane-specific sequencing batch effects we inspected the output of the CollectRnaSeqMetrics tool of Picard Tools version 2.6 [45]. Mapping efficiency and proportion of reads mapping to rRNA, intronic, intergenic and coding regions were obtained from the output of the CollectRnaSeqMetrics (S1 and S2 Figs).

For the poly(A) capture dataset [4], raw FASTQ files were obtained from the Gene Expression Omnibus (GEO:GSE68719) and analyzed exactly as described for our cohorts (with the exception of --rna-strandness in HISAT2, which was turned off to take into account that the cDNA library of this cohort was unstranded).

### RNA expression quantification and filtering

We used Salmon version 0.9.1 [46] to quantify the abundance at the transcript level with the fragment-level GC bias correction option (--gcBias) and the appropriate option for the library type (-l ISR) against the Ensembl release 75 transcriptome. Transcript-level quantification was collapsed onto gene-level quantification using the tximport R package version 1.8.0 [47] according to the gene definitions provided by the same Ensembl release. We filtered out genes in non-canonical chromosomes and scaffolds, and transcripts encoded by the mitochondrial genome. To further reduce the potential for artifacts we filtered out transcripts with unusually high expression by removing transcripts that gathered more than 1% of the reads on more than half of the samples, which resulted in the removal of 3 and 4 transcripts from the PW and NBB cohorts, respectively. Additionally, low-expressed (i.e. genes whose expression was below the median expression in at least 20% of the samples) were filtered out from downstream analyses. Samples were then marked as outliers if their median correlation in gene expression (log counts per million) with the other samples was below Q_1_-1.5*IQR or above Q_3_+1.5*IQR (*Tukey’s fences*; Q_1_: first quartile, Q_3_: third quartile, IQR: inter-quartile range). As a result, 3 samples were marked as outliers in the PW cohort and 3 in the NBB cohort, and were not included in downstream analyses (resulting sample sizes: N_PW_ = 26, N_NBB_ = 18, S4 Fig).

### Estimation of cell composition

For each sample we calculated correlates for the main cortical cell type types (neurons, oligodendroglia, microglia and astrocytes) by summarizing the expression of their respective marker genes as the first principal component of their expression (marker gene profiles, MGPs (Mancarci et al., 2017)). Cortical cell type markers were selected using the homologene R package version 1.5.68 [48]. To unravel potential complex interactions between MGPs and other experimental covariates, including disease status, we calculated the pairwise correlation between all the variables and also their association with the main axes of variation of gene expression. To assist us in choosing an optimal set of MGPs to include as covariates, we quantified the group differences in the cellular proportions between PD and controls using linear models adjusting for the known experimental covariates (i.e. RIN, PMI, sex, age, and sequencing batch). Significant association with disease status was found for oligodendrocyte MGP in the PW cohort and for microglia in the NBB cohort. Thus, these were included in the downstream analyses.

### Differential gene expression and functional enrichment analyses

We performed differential gene expression analyses using the DESeq2 R package version 1.22.2 [49] with default parameters. Experimental covariates (sex, age, RIN, PMI, and sequencing batch) as well as oligodendrocyte and microglia MGPs were incorporated into the statistical model. Multiple hypothesis testing was performed with the default automatic filtering of DESeq2 followed by FDR calculation by the Benjamini-Hochberg procedure. Analyses were carried out independently for the two cohorts. Genes were scored according to their significance by transforming the p-values to account for direction of change. For each gene, the up-regulated score was calculated as *S*_up_ = 0.5 + sign(LFC) · *p*/2, and the down-regulated score as *S*_down_ = 1 − *S_up_*, where LFC corresponds to the log fold change and *p* to the nominal p-value of the gene. Genes were then tested for enrichment using alternatively *S*_up_ and *S*_down_ scores employing the gene score resampling method implemented in the ermineR package version 1.0.1 [50], an R wrapper package for ermineJ [51] with the complete Gene Ontology (GO) database annotation [52] to obtain lists of up- and down-regulated pathways for each cohort.

In order to characterize the main biological processes affected by the cell type correction, we scored pathways based on the loss of significance caused by the addition of cellular estimates to the gene expression model. We quantified the difference in the level of significance in the up- and down-regulated enrichment results for each significant pathway as Δ= log(*p*_0_) − log(*p_CT_*), where *p_CT_* and *p*_0_ are the enrichment p-values for the model with cell types (CT) and without (0), respectively. Only pathways that were significant in either one of the models were analyzed in this manner (*p*_0_ < 0.05 or *p_CT_* < 0.05).

The source code for the analyses is available in the GitLab repository (https://git.app.uib.no/neuromics/cell-composition-rna-pd) under the GPL public license v3.0.

## Supporting information

S1 Figure. Read mapping efficiency

S2 Figure. Read mapping statistics

S3 Figure. Cellular estimates grouped by status

S4 Figure. Sample clustering

S1 File. Significant genes overlapping PW and NBB

S2 File. Complete results of the differential expression analyses

S3 File. ErmineR pathway enrichment analyses before and after correcting for cell composition

S4 File. Up- and down-regulated pathways ranked by the change in p-value resulting from correcting for cell composition

S5 File. Cohort demographic and experimental information

S6 File. Read count matrix

## Supporting information

**S1 Figure. Read mapping efficiency.** (A) Proportion of reads uniquely aligned to the genome. (B) Counts per million (CPMs) for rRNA sequences (PW: ParkWest, NBB: Netherlands Brain Bank, PA: poly(A) capture cohort). Asterisks indicate significance at (*) p > 0.05, (**) p ≤ 0.01, (***) p ≤ 0.001, (****) p ≤ 0.0001, Wilcoxon test.

**S2 Figure. Read mapping statistics.** (A) Percentage of bases mapping to different genomic regions for the two cohorts analyzed using rRNA depletion (PW: ParkWest, NBB: Netherlands Brain Bank) and the poly(A) selection cohort (PA). (B) Scatterplots of samples show RIN values versus percent of bases (unaligned, intergenic, intronic and mRNA). Linear regression trend represented by a black line. Only significant linear regression p-values (p < 0.05) are shown in the panels (PW: ParkWest, NBB: Netherlands Brain Bank, PA: poly(A) capture kit). Asterisks indicate significance at (*) p > 0.05, (**) p ≤ 0.01, (***) p ≤ 0.001, (****) p ≤ 0.0001, Wilcoxon test.

**S3 Figure. Cellular estimates grouped by status.** Astrocyte and neuronal MGPs adjusted for all the experimental variables (sex, age, PMI, RIN, and sequencing batch) and oligodendrocyte and microglia MGPs, but not for disease status. PW: ParkWest, NBB: Netherlands Brain Bank. P-values calculated with Wilcoxon test.

**S4 Figure. Sample clustering.** Samples were marked as outliers if the median correlation with the other samples was below Q1-1.5*IQR or above Q3+1.5*IQR, where IQR stands for inter-quartile range. For each of the rRNA depletion cohorts, the heatmaps represent the pairwise correlation coefficients between the samples based on the log CPM. Marker gene profile (MGP) estimates for the main cortical cell types are represented on the top of each heatmap, together with RIN, disease status, and the outlier status for each sample (PW: ParkWest, NBB: Netherlands Brain Bank).

**S1 File. Significant genes overlapping PW and NBB.**

**S2 File. Complete results of the differential expression analyses.**

**S3 File. ErmineR pathway enrichment analyses before and after correcting for cell composition.**

**S4 File. Up- and down-regulated pathways ranked by the change in p-value resulting from correcting for cell composition.**

**S5 File. Cohort demographic and experimental information.**

**S6 File. Read count matrix.**

## Ethics declarations

### Availability of data and materials

The datasets supporting the conclusions of this article are included within the article and its supplementary files. The source code for the analyses is available in the GitLab repository (https://git.app.uib.no/neuromics/cell-composition-rna-pd) under the GPL public license v3.0.

### Author contributions

GSN participated in the study conception, preprocessed the RNA-seq data, designed and performed the RNA-seq computational analyses. FD participated in the RNA-seq computational analyses. KH contributed to data interpretation and critical revision of the manuscript. IJ and KP participated in the conception and design of the study, data interpretation and critical revision of the manuscript. OBT and GA contributed part of the tissue material, supplementary data, and critical revision of the manuscript. CT conceived, designed and directed the study, contributed to data analyses and interpretation and acquired funding for the study. GSN and CT wrote the manuscript with the active input and participation of LT, KH, IJ, and KP All authors have read and approved the manuscript.

