## Supplementary figures and images for "Common gene expression signatures in Parkinson’s disease are driven by changes in cell composition"

### S1 Figure. Read mapping efficiency

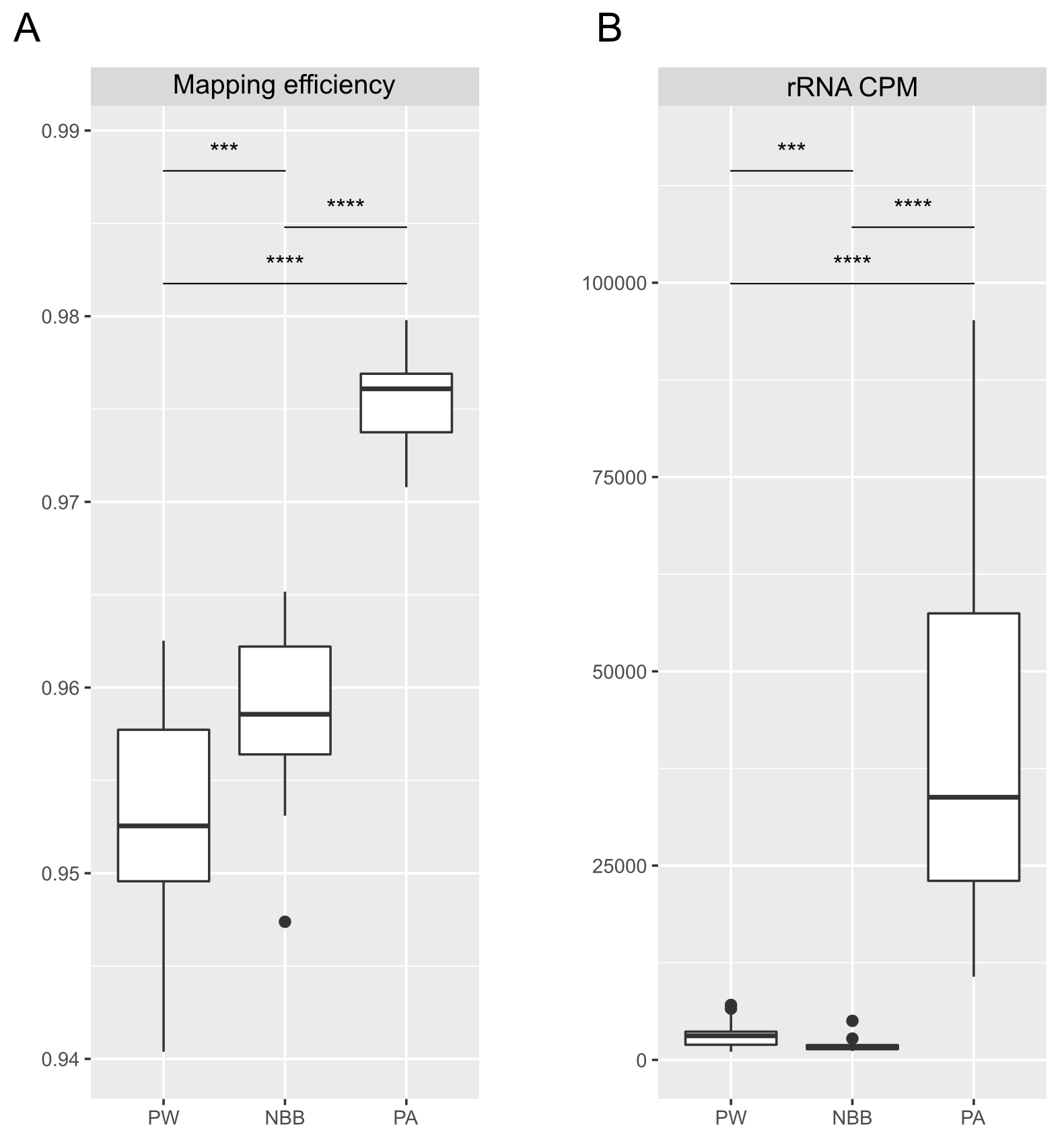

### S2 Figure. Read mapping statistics

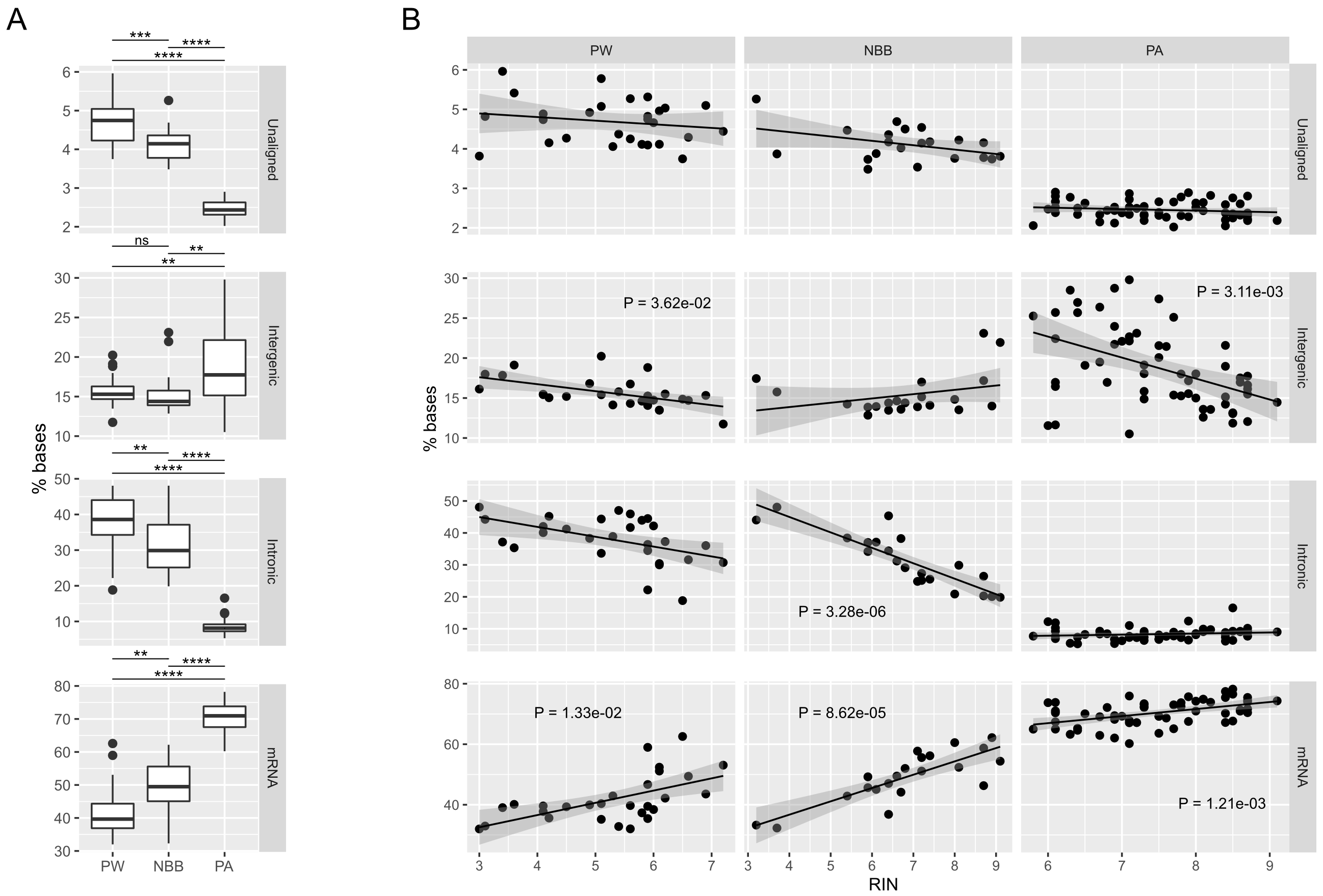

### S3 Figure. Cellular estimates grouped by status

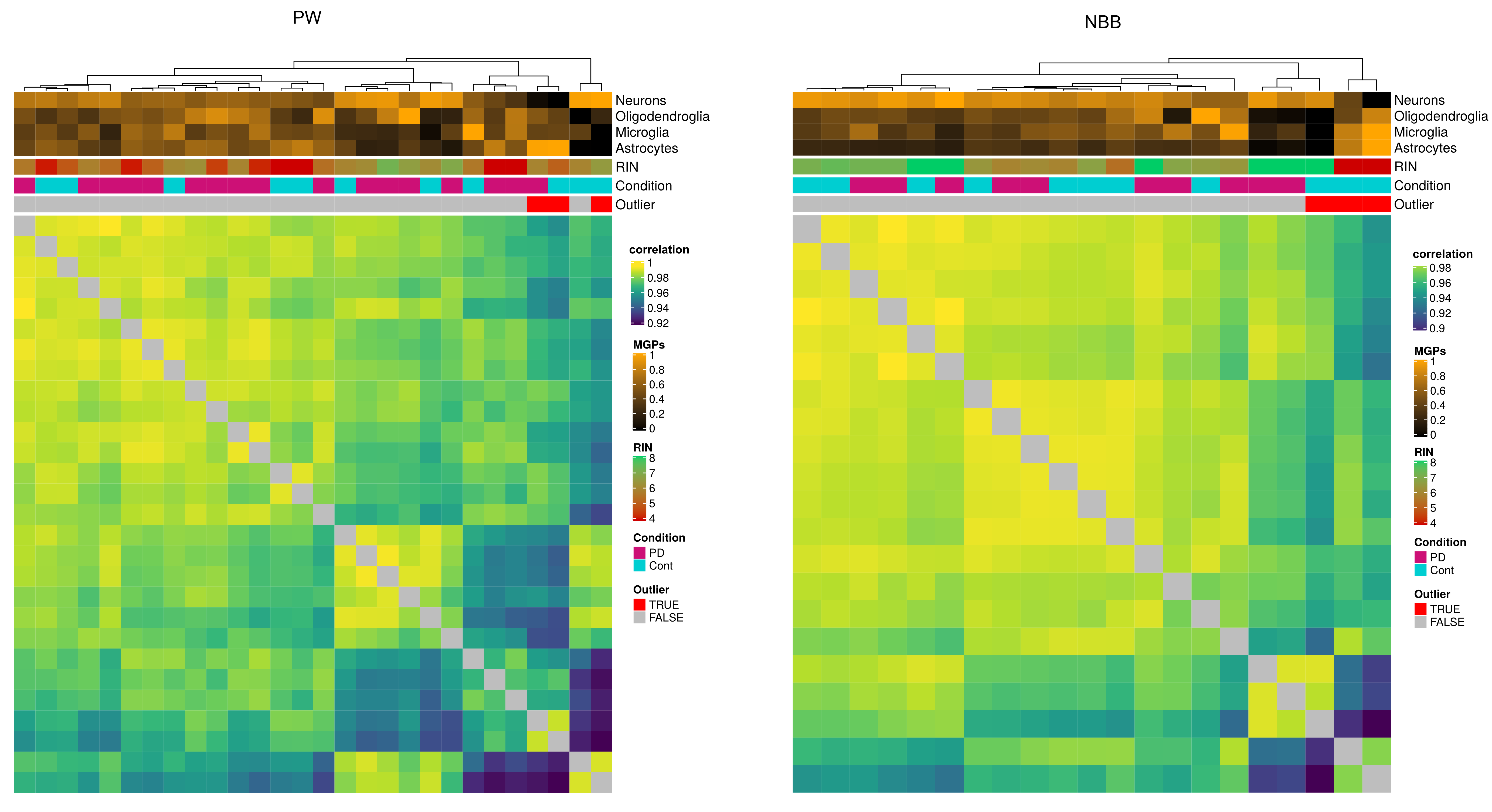

### S4 Figure. Sample clustering

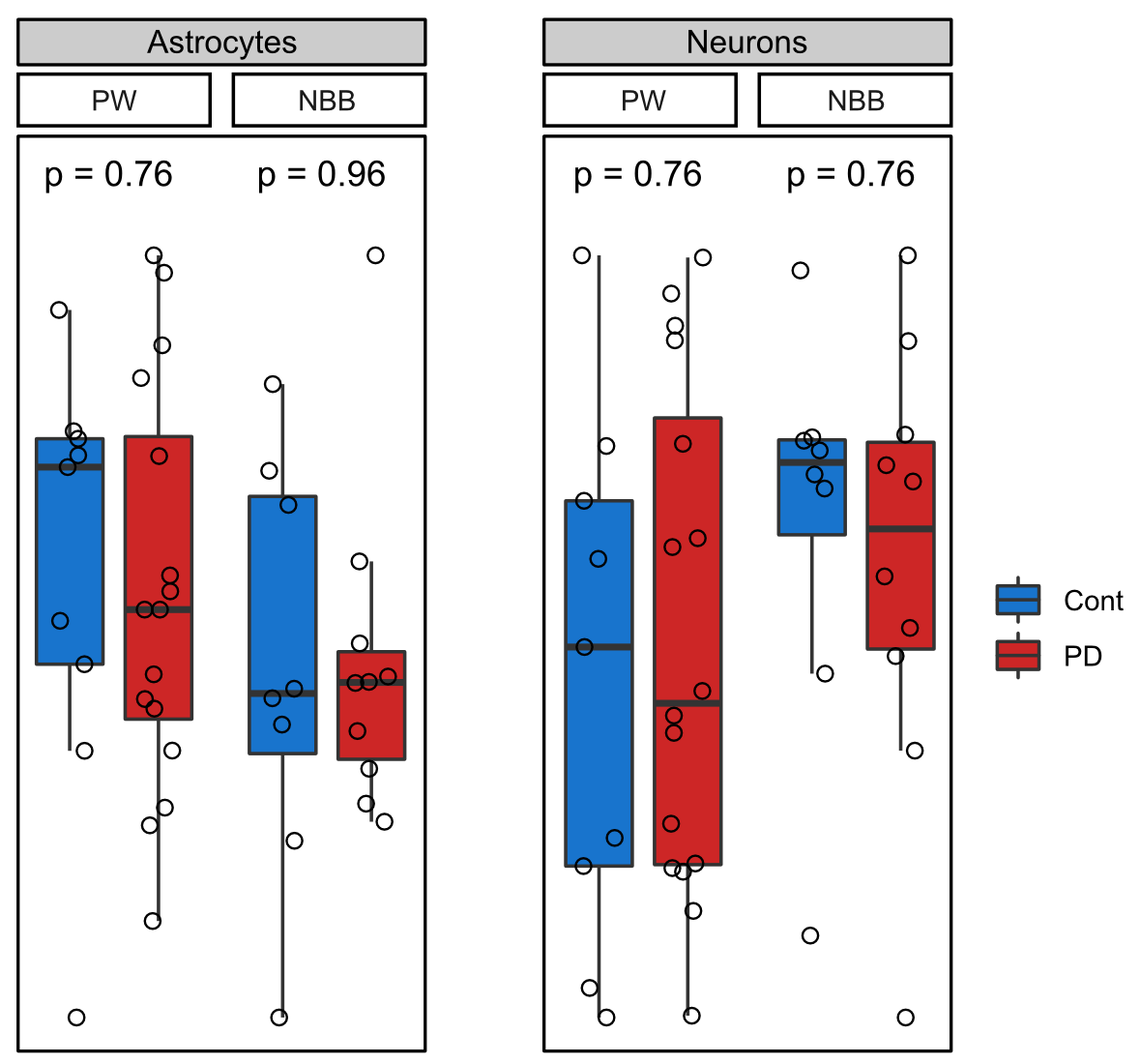
